## Supplementary information for "Unmasking AlphaFold: integration of experiments and predictions in multimeric complexes"

May 13, 2024

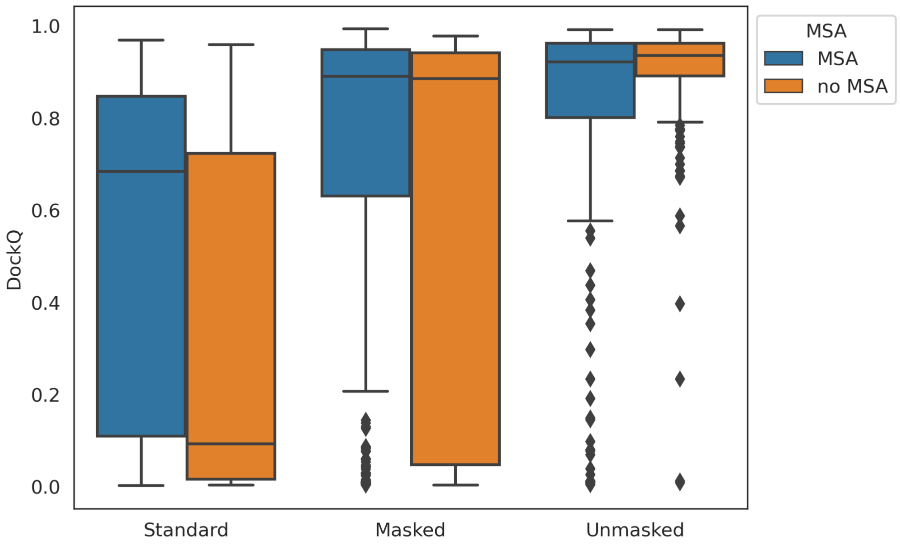


Figure S1: Box plot comparison of various template strategies when predicting a subset of the PDB containing heterodimer complexes. These targets were selected from structures deposited in the PDB after the parameters of the latest version of AlphaFold-Multimer was released. The default template strategy has been followed in the tests, first by letting AlphaFold perform its own template search on an older PDB ("Standard") and also by making AlphaFold use the native structures themselves ("Masked"). These are compared with AF_unmasked when the cross-chain constraints are included in the prediction step and the native structures are used as templates ("Unmasked"). Masked and Unmasked predictions are done both with regular MSA inputs ("MSA") and by completely clipping MSAs so that only the target sequences are included ("no MSA"). We use DockQ to score the top-ranked prediction by AlphaFold's predicted ranking confidence. Unmasked predictions are better in both scenarios, but they get better when MSA inputs are clipped. Since we are using ideal templates, Masked predictions tend to be better than Standard predictions even when cross-chain information is not used.

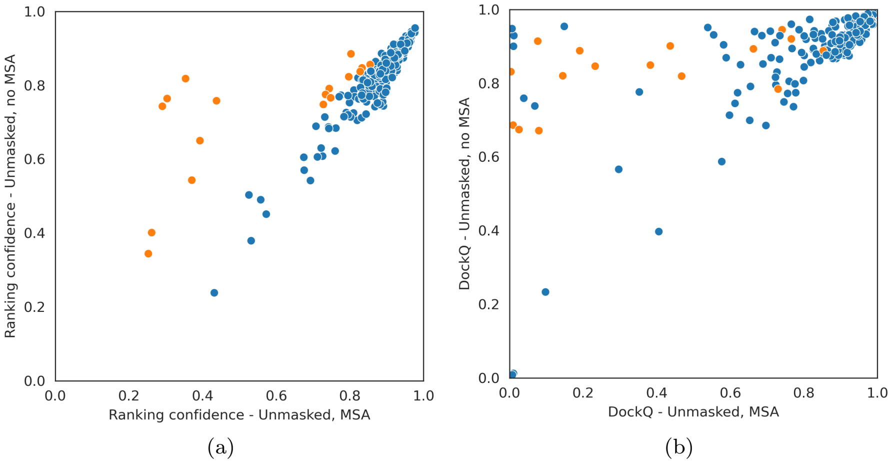


Figure S2: Scatter plot comparison of Unmasked predictions when using (”MSA”) or clipping evolutionary inputs (”no MSA”) to only include the target sequence. We use DockQ to score the top-ranked prediction by AlphaFold’s predicted ranking confidence. While ranking confidences correlate, the overall confidence in most cases is lower when no MSA inputs are included (a, blue). But in the cases where the ranking confidence actually increases when clipping the MSA (a, orange), predictions are also better (b, orange).

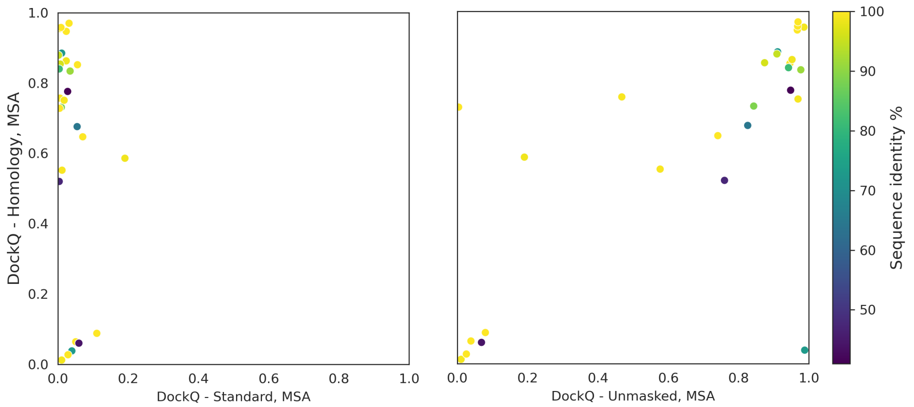


(a) (b)

Figure S3: (a) Scatter plot comparison of Standard AlphaFold predictions and predictions where good homology models were used in AF_unmasked as templates. (b) Scatter plot comparison of AF_unmasked predictions where native structures were used as template and AF_unmasked predictions where good homology models were used as templates. The similarity between the homologous template and the target sequences (measured by sequence identity) does not seem to correlate with the quality of the predictions.

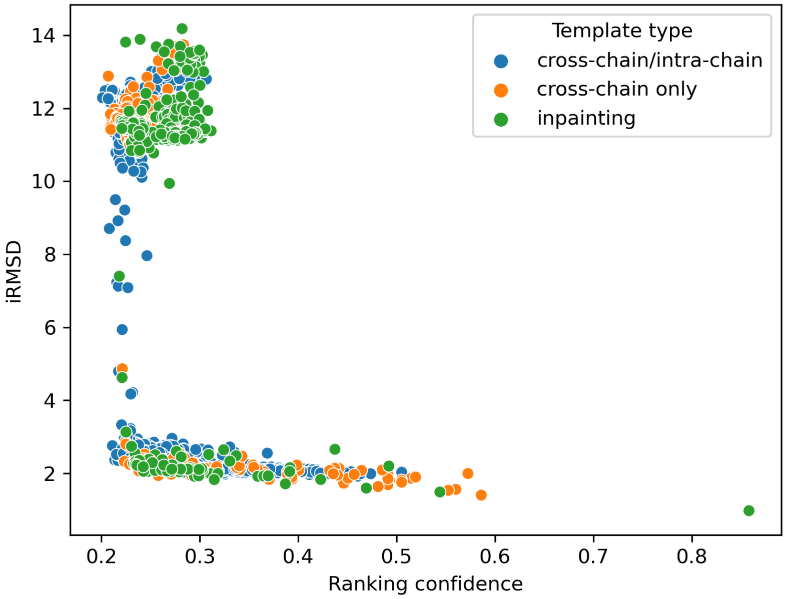


Figure S4: Scatter plot of AlphaFold ranking confidence and interfacial residue RMSD (iRMSD) for all AF_unmasked predictions for CASP15 target H1142. The models with better quality (low iRMSD) have ranking confidence that correlates very well with iRMSD. The top ranked model (bottom-right) is also the model with lowest iRMSD.

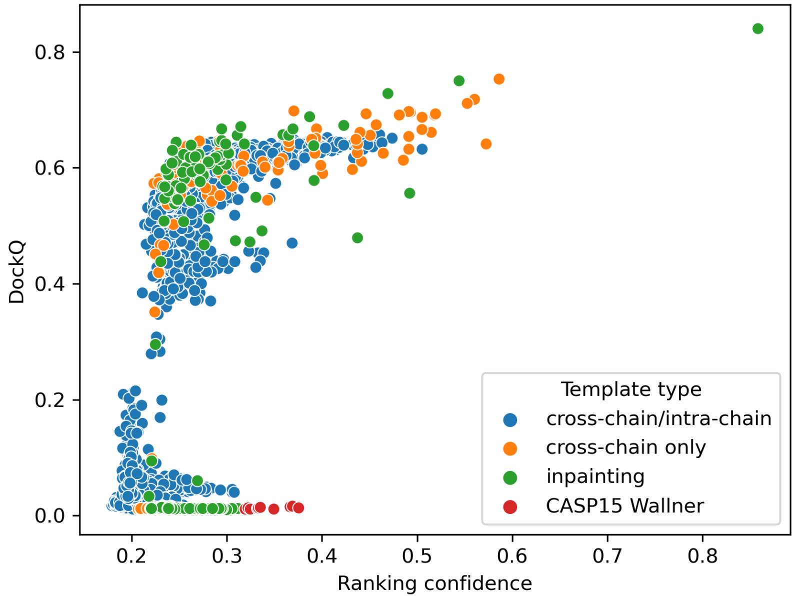


Figure S5: Scatter plot of AlphaFold ranking confidence and DockQ score for all AF_unmasked predictions for CASP15 target H1142. The ranking confidence predicted by AlphaFold correlates well with the quality of the model as calculated by DockQ. The top ranked model (top-right) is also the model with highest DockQ score.

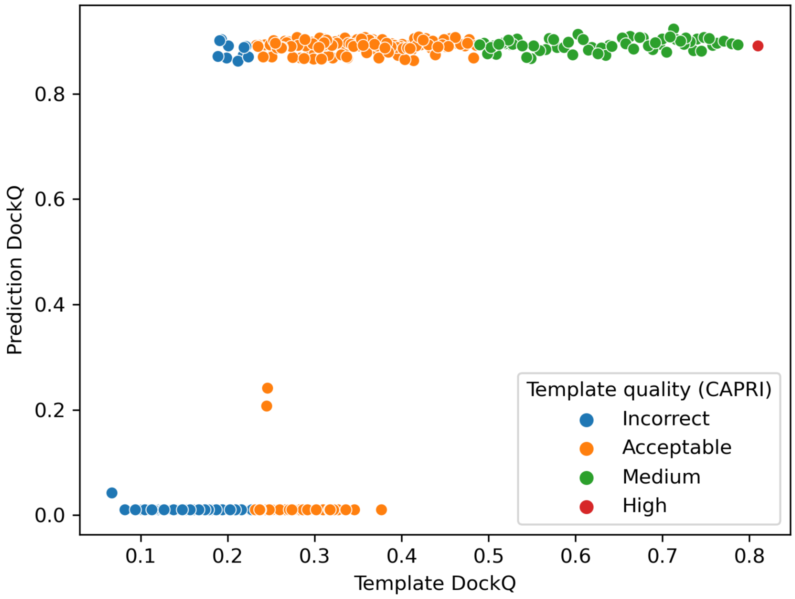


(a)

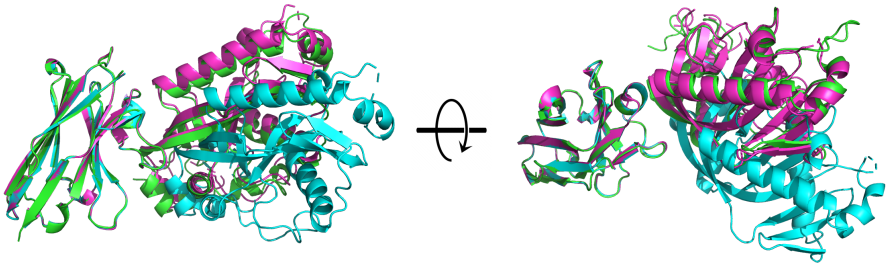

(b)

Figure S6: (a) Scatter plot of template and prediction DockQ scores for target H1142 when the native is randomly perturbed 300 times around the native conformation with RosettaDock. The top-ranked (by AlphaFold's ranking confidence) prediction starting from each perturbed template is evaluated with DockQ and compared to the quality of the initial template. Each point is colored by template correctedness as defined in the CAPRI community. (b) Example of prediction where AF_unmasked was able to recover the correct conformation from an imperfect template. The native is shown in magenta, the initial template in cyan (DockQ score: 0.19) and the top-ranked prediction in green (DockQ score: 0.87). All structures are superimposed by the ligand chain.

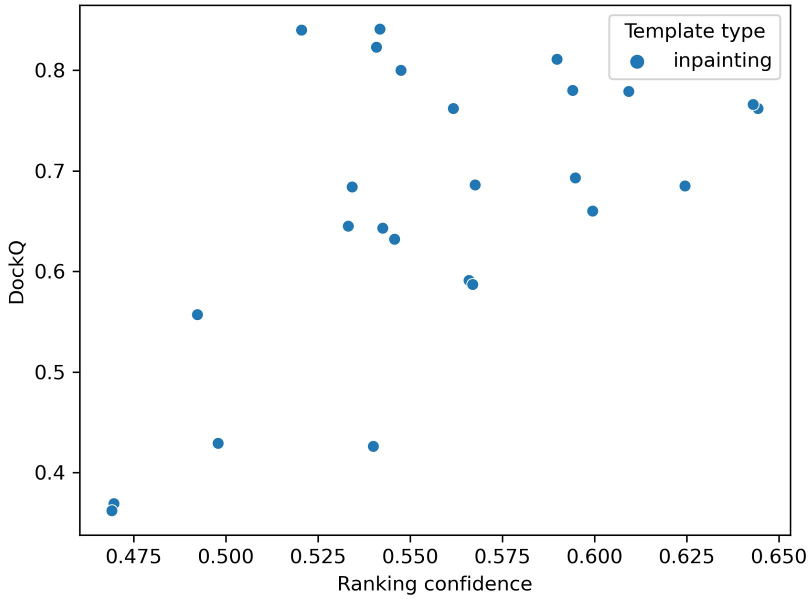


Figure S7: Scatter plot of AlphaFold's ranking confidence and DockQ score for CASP15 target H1111. Not all predicted areas here are evaluated, as large domains that were inpainted by AF_unmasked could not be evaluated against the native. The top 3 ranked models (right side) show a variety of potentially biologically relevant conformations in the inpainted areas.

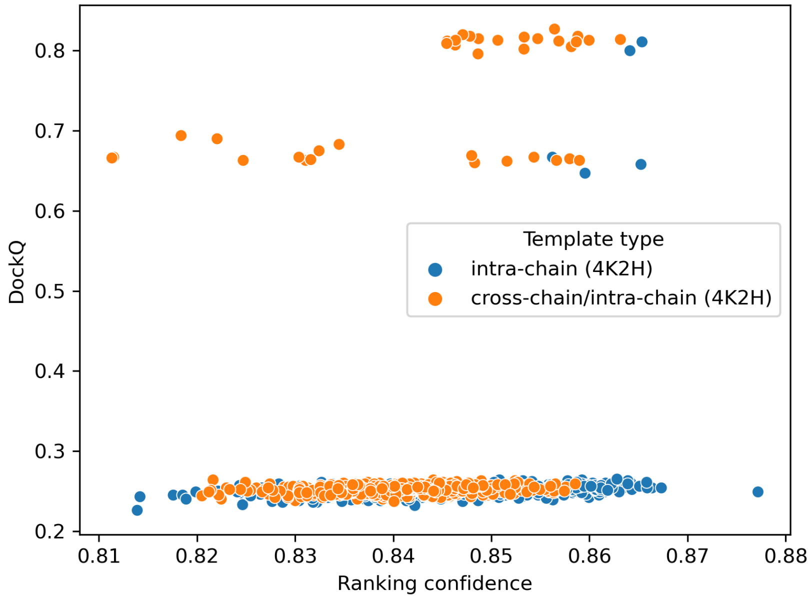


Figure S8: Scatter plot of AlphaFold's ranking confidence and DockQ score for CASP15 target T1109o.

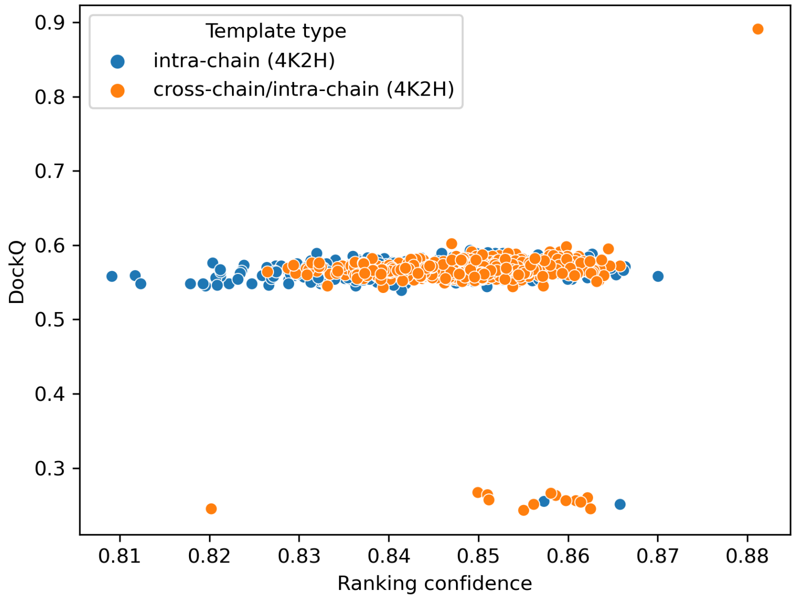


Figure S9: Scatter plot of AlphaFold's ranking confidence and DockQ score for CASP15 target T1110o.

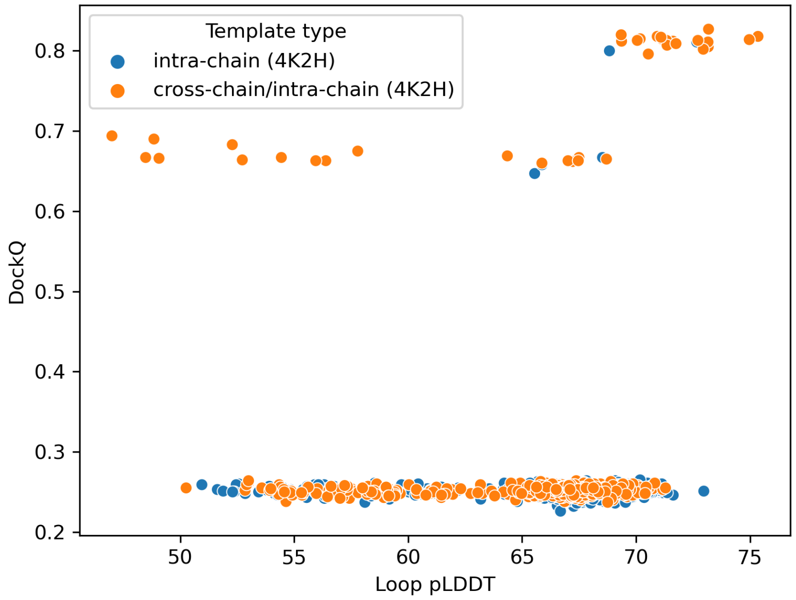


Figure S10: Scatter plot of AlphaFold's predicted LDDT score (pLDDT) in the inpainted loop areas and DockQ score for CASP15 target T1109o. Here, the pLDDT of the inpainted loops are a better way of separating good models from bad, as local changes in inpainted regions can be fairly small when compared to the prediction as a whole.

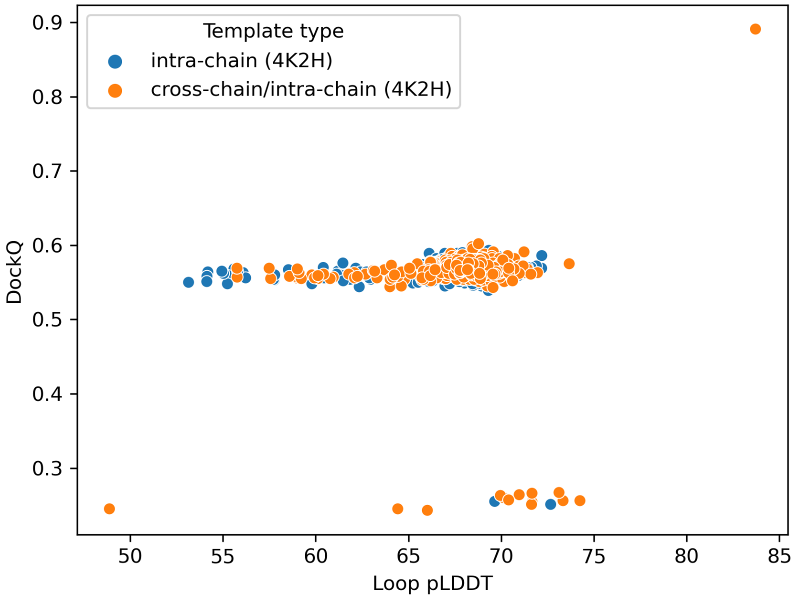


Figure S11: Scatter plot of AlphaFold's predicted LDDT score (pLDDT) for inpainted loop areas and DockQ score for CASP15 target T1110o.

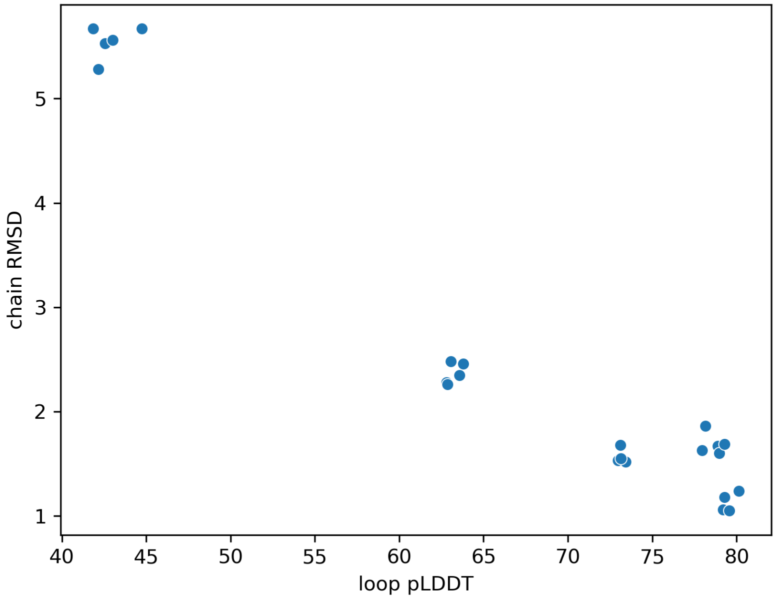


Figure S12: Scatter plot of AlphaFold's predicted LDDT score (pLDDT) for Rubisco's inpainted loop areas and RMSD between predicted small subunit and experimental structure. pLDDT is calculated here as the average pLDDT for $C\alpha$ atoms in the inpainted regions in AF_unmasked predictions. The average is calculated across all eight inpainted loops in each prediction. The RMSD is calculated with TM-align by superimposing all predicted small subunits against all experimental small subunits and picking one of these $8 \times 8$ comparisons that has lowest RMSD.

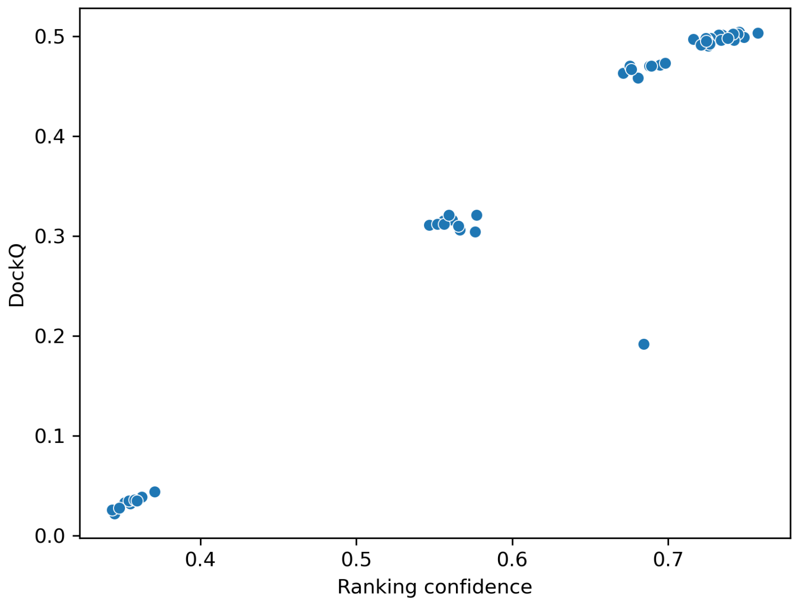


Figure S13: Scatter plot of 50 ClpB hexamer models with inpainting of AAA domains. Inpainted areas are not included in the DockQ calculation.

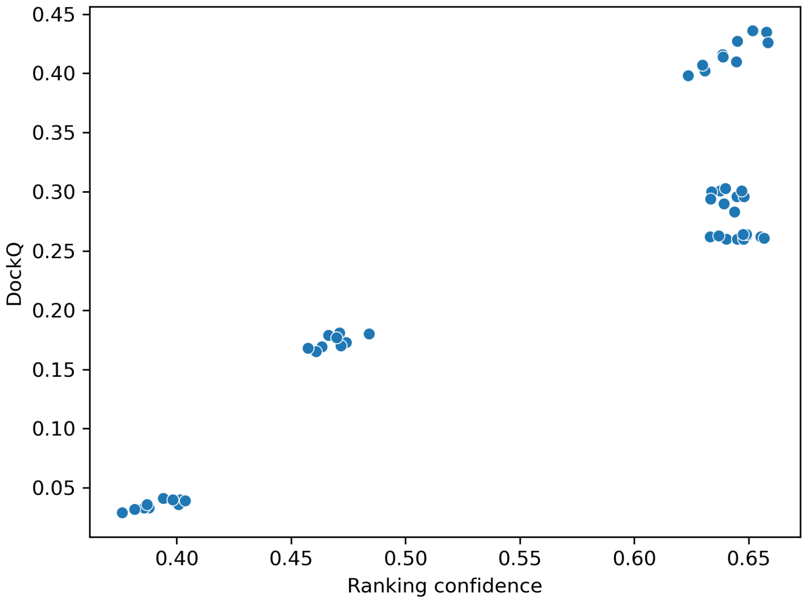


Figure S14: Scatter plot of 50 ClpB hexamer models with inpainting of AAA domains and embedded casein molecule. Inpainted areas are not included in the DockQ calculation.

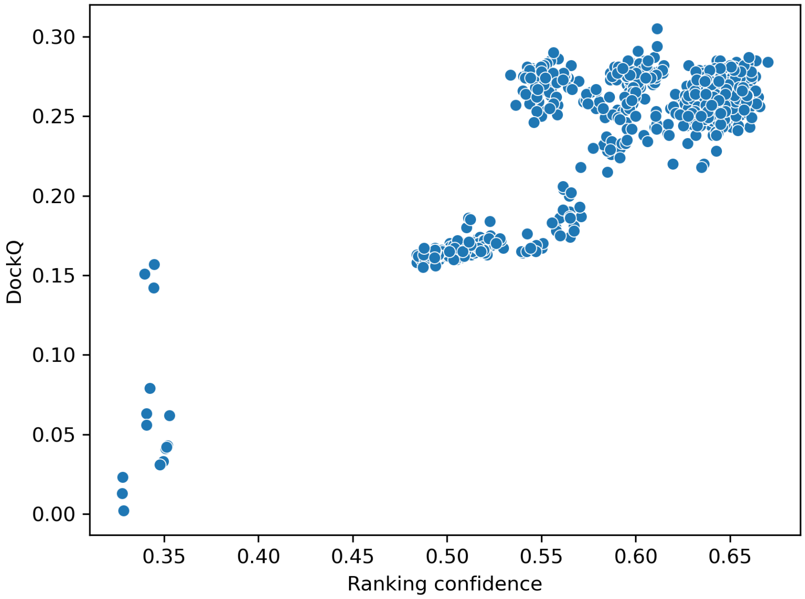


Figure S15: Scatter plot of 1000 NF1 models with inpainting of GRD domains. DockQ is calculated against the deposited structure of NF1 where GRD are in the ”closed” conformation (PDB ID: $7PGU$).

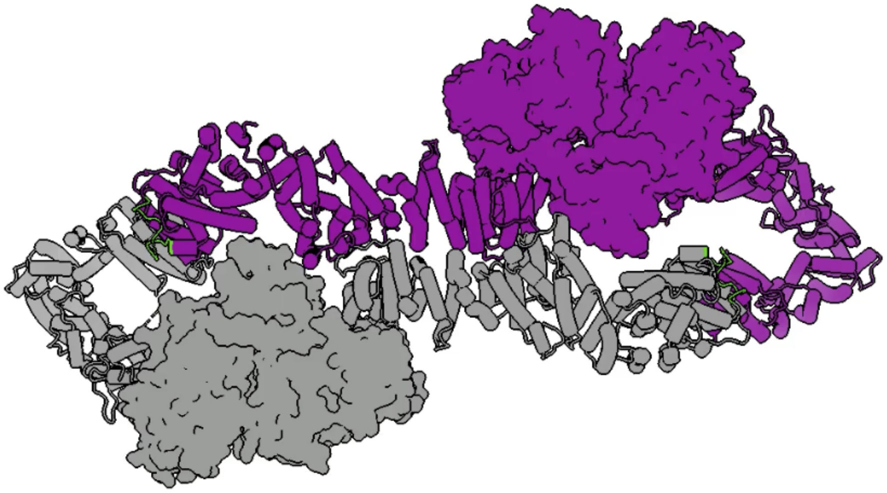

Figure S16: Still image from video of partial opening of NF1. Interpolation between AF_unmasked models showing different positions of the GRD and Sec14-PH regions (shown as surfaces) relative to the helical platform (shown as cartoon). The two chains of NF1 are coloured in grey and purple. Video link: https://figshare.com/s/3f30484e34447d388e90?file=46052964

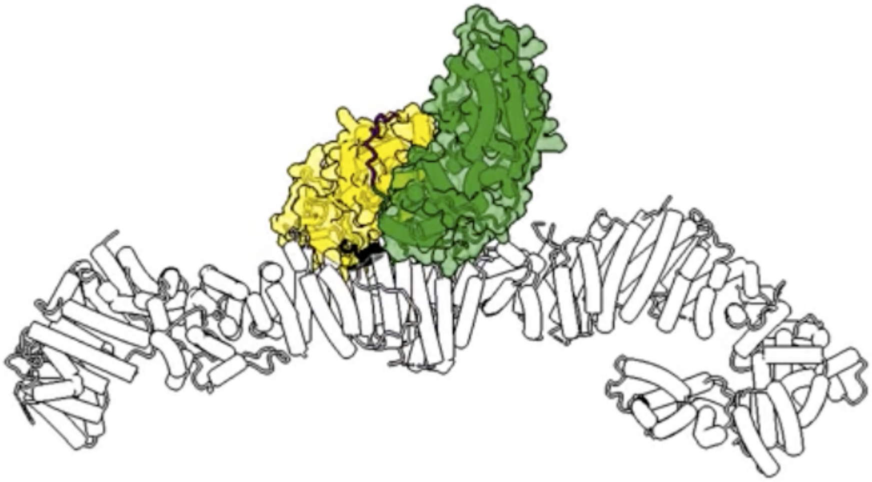

Figure S17: Still image from video of transition from the closed to the open position of NF1. Interpolation between AF_unmasked models from the closed to the open positions of the GRD and Sec14-PH regions, shown in gold and green respectively. Only one monomer is displayed. Video link: https://figshare.com/s/3f30484e34447d388e90?file=46052946

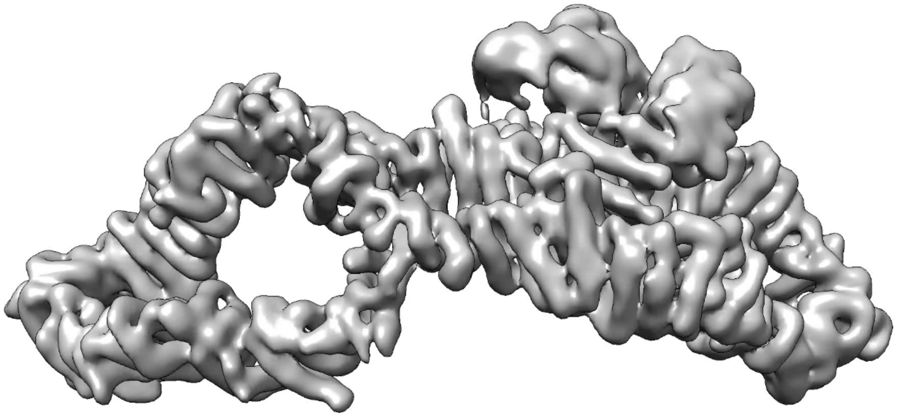

Figure S18: Still image from video of 3D variability analysis of NF1 cryo-EM data, showing bending of the NF1 helical platform as well as appearance and disappearance of the GRD and Sec14-PH regions. Video link: https://figshare.com/s/3f30484e34447d388e90?file=46052958

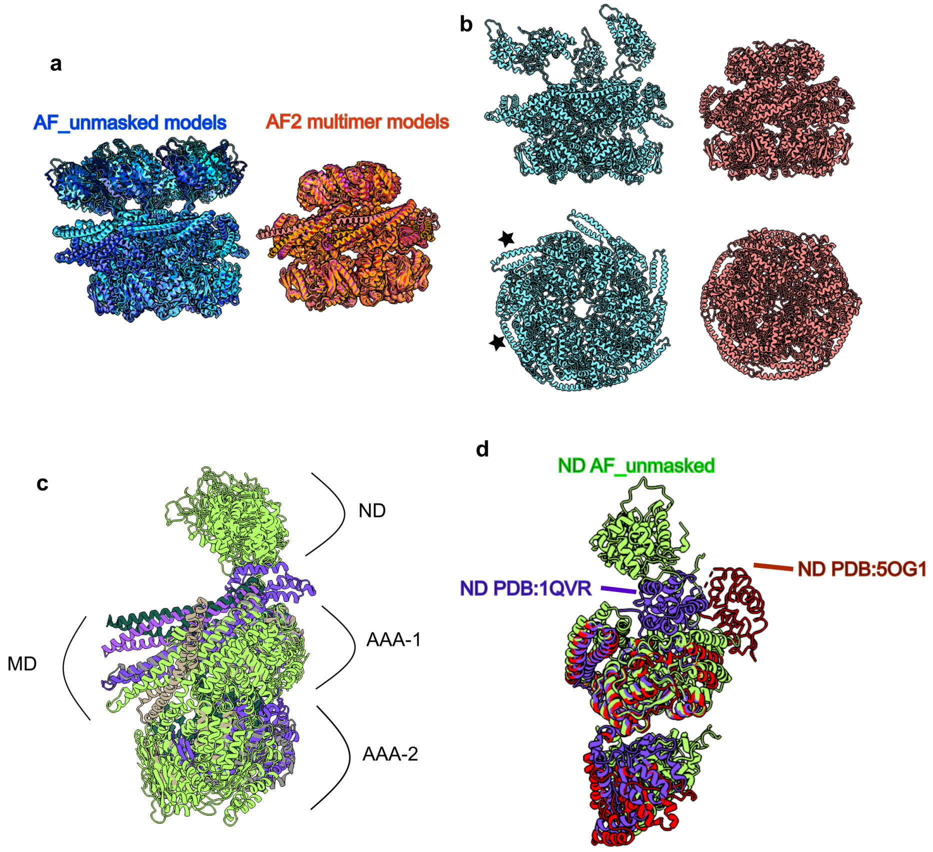


Figure S19: Comparison of the ClpB AF_unmasked predictions with AlphaFold-Multimer (v2.3) predictions and experimental PDB structures. (a) Superimposition of 9 predictions of the ClpB hexamer, performed with either AF_unmasked (shades of blue) or AlphaFold-Multimer (shades of red). The AlphaFold-Multimer predictions are all remarkably symmetric and very compact.(b) Comparison of one ClpB hexamer obtained with AF_unmasked (cyan) and one with AlphaFold-Multimer (salmon).

While AF_unmasked picks different orientations of the M-domain (MD) within the same hexamer (labelled with stars), AlphaFold-Multimer fails to do it. (c) Monomers of ClpB obtained with AF_unmasked (lime green), and deposited structures of ClpB monomers obtained using X-ray diffraction or cryo-EM (various colours) are superimposed on their AAA-1 domain. The MD adopts different orientations as well as the AAA-2 (PDB ID: $6W6E$, $6W6H$, $1QVR$, $4HSE$, $4CIU$, $5OFO$). This shows that the AF_unmasked predictions are compatible with the current structural knowledge of the system. (c) Comparison of the positions of the N-terminal domains of ClpB (ND) as known in experimental data and on the AF_unmasked predictions. Note that the N-terminal domains are linked to ClpB body by long flexible loops.
